# Benchmarking differential abundance methods for finding condition-specific prototypical cells in multi-sample single-cell datasets

**DOI:** 10.1101/2023.02.24.529894

**Authors:** Haidong Yi, Alec Plotkin, Natalie Stanley

## Abstract

Modern single-cell data analysis relies on statistical testing (e.g. differential expression testing) to identify genes or proteins that are up-or down-regulated in relation to cell-types or clinical outcomes. However, existing algorithms for such statistical testing are often limited by technical noise and cellular heterogeneity, which lead to false-positive results. To constrain the analysis to a compact and phenotype-related cell population, differential abundance (DA) testing methods were employed to identify subgroups of cells whose abundance changed significantly in response to disease progression, or experimental perturbation. Despite the effectiveness of DA testing algorithms of identifying critical cell-states, there are no systematic benchmarking or comparative studies to compare their usages in practice. Herein, we performed the first comprehensive benchmarking study to objectively evaluate and compare the benefits and potential downsides of current state-of-the-art DA testing methods. We benchmarked six DA testing methods on several practical tasks, using both synthetic and real single-cell datasets. The task evaluated include, recognizing true DA subpopulations, appropriate handing of batch effects, runtime efficiency, and hyperparameter usability and robustness. Based on various evaluation results, this paper gives dataset-specific suggestions for the usage of DA testing methods.

## Introduction

Modern single-cell technologies have enabled the measurement of thousands of genes of tens of proteins in samples collected in a variety of states, such as development [1, 2, 3], disease progression [4, 5, 6, 7], or after experimental perturbation [8, 9, 10]. In a single-cell dataset collected from multiple patient samples, heterogeneity is always present to some degree among all cell populations. For instance, (i) the number of profiled cells may vary across samples (ii) some cells in a single-cell sample might not respond to experimental perturbations or simply act as background cells in terms of clinical consequences [10]. Such cellular heterogeneity causes true biologically driven signals to be obscured by unrelated variability, making it difficult for downstream analysis to identify them or producing false positive results. Therefore, identifying some “clean” cell populations purely perturbed by the corresponding experimental conditions in a single-cell dataset becomes crucial for further statistical analysis [11, 12].

To tackle this problem, differential abundance (DA) testing methods have been utilized to identify a cohesive subset of cells linked with clinical outcomes of interest (See Figure 1 for an intuitive illustration). DA testing methods accomplish this by identifying regions enriched with cells as a result of biological perturbations. One class of DA testing methods [13, 14, 15] identifies phenotype-associated subgroups of cells by determining if statistically significant changes in abundance occur in response to a biological perturbation. This method can effectively eliminate the cells that are unaffected by treatment condition since they are evenly dispersed across the treatments. Another category of DA testing approaches [10, 16] uses a different approach with no statistical testing.

**Figure 1:**
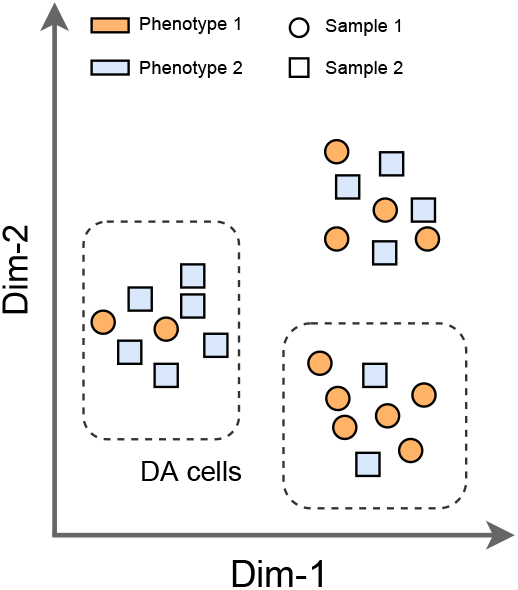
An illustration of DA cells in a dataset containing two samples under two phenotypes.

In contrast, these approaches perform conditional density estimation on cells from various experimental conditions and select phenotypically significant subsets of cells based on their predicted density scores for each condition. DA testing methods can also be divided into two types based on their use of clustering: clustering-based methods [17, 18, 19] and clustering-free methods [10, 13, 14, 15, 16]. Since most clustering algorithms are unstable due to their nonconvexity and can only provide a rough partition for distinct cell states, many clustering-free methods have proven to be preferable to clustering-based methods [10, 13, 14].

Recent single-cell studies have demonstrated considerable success using DA testing methods to identify novel cell states from a broad landscape of profiled single cells. DA testing has been used to reveal enrichment of granulocytes, monocytes, and B cells in patients who died from COVID-19 [20]; identify rheumatoid arthritis (RA)-associated cell populations from a single-cell dataset of 18 patients with either rheumatoid or osteoarthritis [15]; and discover a new subpopulation of cells in mouse intertypical thymic epithelial cells (TECs) that are depleted with age [14]. Despite the fact that numerous research efforts have gone into developing new DA methods, there have been considerably few studies providing thorough and quantitative comparisons of the strengths and weaknesses of the common DA testing approaches, especially those that are clustering-free. Furthermore, in the original papers introducing each new method, most of the results focused on unraveling insights into complicated biological processes and focused on elucidating superiority over clustering-based methods. Among the original DA testing studies, Ref. [14] is the only work that quantitatively compared present state-of-the-art DA testing methods for identifying DA cell populations. However, Ref. [14] presented simply numerical results without additional analysis, restricting their intepretability and comprehension. To address this deficiency, we intend to study current DA testing methods and assess their strengths and weaknesses in various circumstances.

In this benchmarking study, we evaluated six DA testing methods, including both the clustering-based and clustering-free approaches. In our experiments, we compared the six DA testing methods using synthetic and real-world single-cell datasets. We examined various facets of the DA testing methods, such as (1) the precision for detecting DA subpopulations in data with diverse differential trajectory structures; (2) the capacity to handle technical and biological variables, such as batch effects; (3) runtime efficiency and scalability; and (4) usability and robustness with regard to hyperparameters. To aid in a better understanding, as a key result, we established in a synthetic dataset that several DA methods cannot perform well when the number of cells is significantly unbalanced between DA subpopulations. After investigating the characteristics of each method in relation to the unique characteristic of each of the diverse datasets, we ultimately provided data-specific suggestions for choosing the best DA testing methods to use in clinical settings.

## Main

### Description of datasets

In this study, we evaluated the performance of six prevalent DA testing methods on three simulated datasets, a single-cell RNA sequencing dataset (scRNA-seq), COVID-19 PBMC [5] and a CyTOF dataset, BCR-XL [8]. To facilitate thorough benchmarking, the experimental datasets differ in several ways, including, topology of differential trajectories, ratio or *extent* of differential abundance (DA ratio), and single-cell modality (e.g. protein vs gene measurement). The three synthetic datasets, for example, have different topological structures (linear, branch, and cluster) of their differential trajectories and DA ratios. The COVID-19 PBMC dataset measures the expression of genes, whereas the BCR-XL dataset measures the expression of proteins.

### Synthetic Datasets

Using the R package dyntoy (described in Ref. [14]), we generated count matrices for three synthetic single-cell datasets with diverse topological structures, including linear trajectories, branching trajectories, and discrete clusters. Each dataset included six samples from a simulated experiment with three replicates (R1, R2, and R3) and two experimental conditions (C1 and C2). Different datasets may contain distinct cell populations of differing sizes. For example, the linear and branch datasets contain 7500 cells and 500 genes, whereas the cluster dataset contains 2,700 cells and 500 genes. To quantitatively benchmark DA testing methods, we generated ground-truth DA labels as follows: First, we selected one of the cell populations as the target population to exhibit differential abundance between the two experimental conditions, while considering the remaining cell populations as background cells. Next, using the same pipeline provided in Ref. [14], we calculated the probability for each cell under each condition (*P*(*C*1) or *P*(*C*2)). Lastly, we randomly assigned the ground-truth labels to each cell based on the conditional probability. To facilitate a thorough evaluation, for each dataset, we simulated different DA labels by varying (1) the target DA cell population; (2) the DA ratio in the target cell population; and (3) the random seed. Furthermore, we also simulated batch effects with varying magnitudes in the synthetic dataset by adding noises sampled from isotropic normal distribution with different variance. It is worth noting that in Ref. [14], the ground truth DA label for each cell is determined by a constant percentile threshold, *t* of the distribution *P*(*C*2) across simulations. However, in our strategy, *t* is adjusted adaptively depending on the size of the target cell population throughout the entire dataset. Formally, our *t* threshold is defined as,

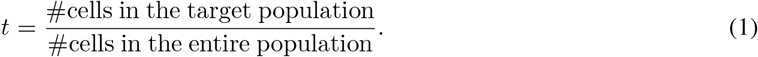

Here, # represents the ‘number’, or count of cells in the respective bins of target or entire population. Visualizations for each of the synthetic datasets are shown in Supplementary Figure S1.

### COVID-19 PBMC Dataset

The COVID-19 PBMC dataset is a single-cell RNA-sequencing dataset generated by profiling 44,721 peripheral blood mononuclear cells (PBMCs) from seven hospitalized COVID-19 patients, four of whom had acute respiratory distress syndrome, and six healthy controls [5]. Considering that one of the patients has two replicates (A and B), the COVID-19 dataset contains a total of 14 samples with different clinical symptoms, including 8 COVID-19 samples and 6 healthy controls. In addition to basic clinical outcomes, this dataset contains information regarding the COVID-19 illness course of each patient, such as, severity classification at the time of admission (ICU/Floor), ventilation status, etc. In the original study (Ref. [5]), the authors investigated the changes in cell type proportions between the COVID-19 samples and the healthy controls and revealed that case severity was associated with the depletion or expansion of several canonical immune cell-types, including developing neutrophils and plasmablast. Therefore, the intended use of this dataset was to examine the efficacy of different DA testing approaches for identifying differentially abundant cell-populations.

### BCR-XL CyTOF Dataset

The BCR-XL dataset presented in Ref. [8] is comprised of 172,791 human PBMCs collected across 16 CyTOF samples, eight of which were stimulated with B cell receptor/Fc receptor cross-linker (BCR-XL) with the remaining being case controls. Cells in this dataset belong to one of eight manually-gated cell populations, including B-cells IgM-, B-cells IgM+, CD8+ T-cells, CD4+ T-cells, DC, surface-cells, NK cells, and monocytes. There are 35 measured markers in the BCR-XL dataset, but we only preserved 24 functionally meaningful markers in our experiments.

### Benchmarking overview

In this benchmarking study, we intend to compare current state-of-the-art DA testing methodologies for identifying phenotype-associated cell-populations in an impartial and thorough manner, as well as to investigate their strengths and potential drawbacks in single-cell data analysis. Figure 2 depicts our benchmarking workflow. To assess the performance of DA testing methods for predicting DA cell populations, we used mass cytometry (CyTOF), single-cell RNA sequencing (scRNA-seq), and synthetic datasets created using dyntoy [21] and Splatter [22]. These experimental datasets covered a diverse range of biological contexts. For example, the synthetic datasets had various cell spreading topological structures in high-dimensional space, such as linear, branch and cluster, which each reflected a unique single-cell differential trajectories (see Supplementary Figure S1). We further included diverse, real CyTOF and scRNA-seq datasets with clinical outcomes, including peripheral blood mononuclear cells (PBMCs) profiled in patients with COVID-19 PBMC [5], and a BCR-XL CyTOF dataset measuring human PBMCs stimulated with B cell receptor/Fc receptor cross-linker (BCR-XL) [8].

**Figure 2:**
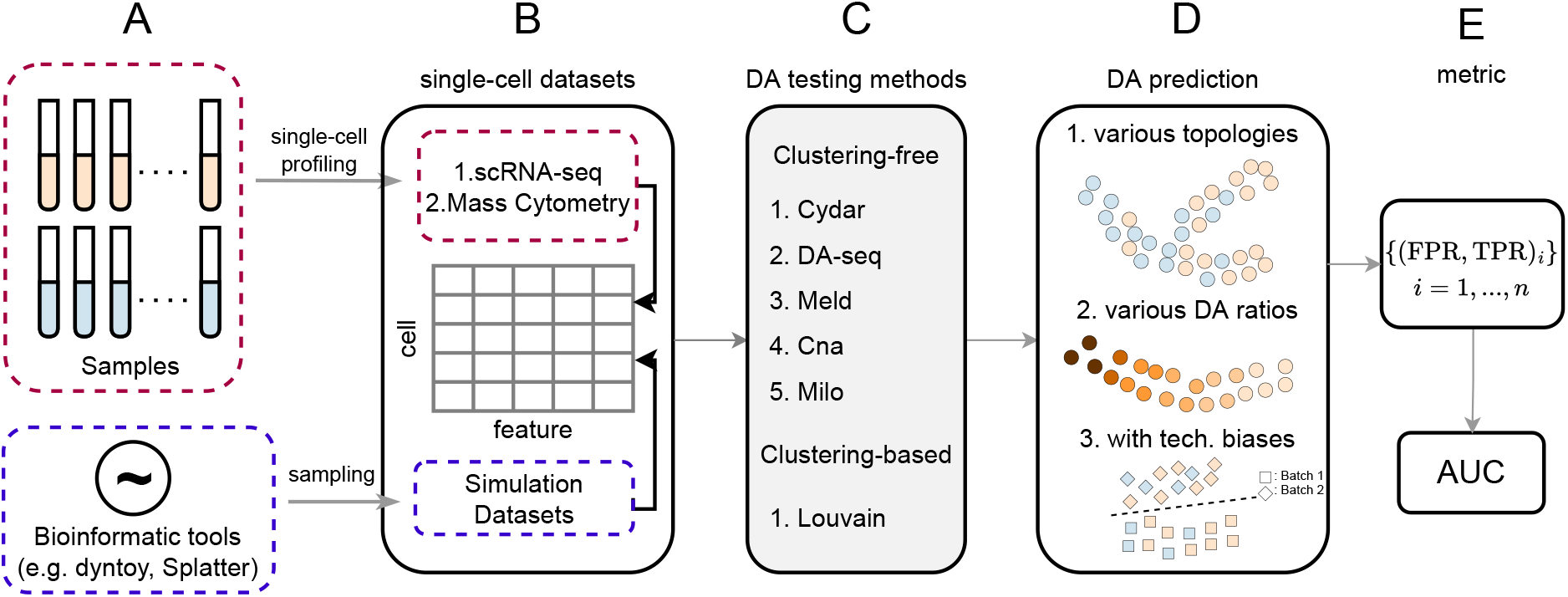
Schematic illustration of the benchmarking workflow. Using both synthetic and real single-cell datasets, six DA testing methods were evaluated under three configurations for the DA prediction task. (A)&(B). First, single-cell RNA-seq and mass cytometry datasets are collected from profiled patients, or a synthetic datasets are generated using the packages dyntoy [21] or splatter [22]; (C)&(D). Next, we evaluated the six clustering-based [23] and clustering-free [13, 16, 10, 14, 15] DA testing methods on datasets with different topologies, DA ratios, and technical biases such as batch effects; (E). Lastly, we compare the performance of the DA testing methods using AUC score.

We evaluated a total of six distinct DA testing approaches, which can be broadly categorized into two groups: (i) clustering-based methods [23], and (ii) clustering-free methods [13, 16, 10, 14, 15]. For clustering-free methods, we benchmarked five methods including Cydar [13], Milo [14], DA-seq [16], Meld [10] and Cna [15]. Noting that clustering-free methods often exhibit superior performance in comparison to clustering-based methods, we compared all such results to the Louvain algorithm [23], a commonly used graph-based clustering method in single-cell data analysis. To further show the differences of the six DA testing methods, we compare their properties in Supplementary Table S1. Here, we provide a brief summary of the six DA testing approaches included in the following benchmark study. For more implementation details, see section Differential abundance (DA) testing methods in Methods.

1. Cydar [13]: Cydar detects DA cell populations by assigning cells to hyperspheres and testing whether the number of cells in each hypersphere varies in a statistically-significant way between conditions. The spatial false discovery rate (FDR) throughout the high-dimensional space controls Cydar’s Type I error;
2. DA-seq [16]: DA-seq predicts DA scores for each cell under two separate conditions by applying a logistic regression model. Label permutation is then used to empirically evaluate the statistical significance of the prediction results;
3. Meld [10]: Meld calculates the likelihood that each cell belongs to or is prototypical of each condition, using a graph-based kernel density estimation (KDE) method. The DA cells are then selected by setting a heuristic likelihood threshold;
4. Cna [15]: Cna uses random walks on graphs to generate a neighborhood abundance matrix (NAM), which quantifies the relative abundance of each sample within particular cellular neighborhoods. DA cell-populations are then ultimately identified through statistical testing based on the NAM across the conditions.
5. Milo [14]: Milo begins by counting the number of cells of each sample within k-nearest neighborhoods and then applies a negative binomial generalized linear models (NB-GLM) to test the DA of each local graph. Milo, like Cydar, controls type-I error via spatial FDR;
6. Louvain [23]: The Louvain method first clusters cells across samples using the Louvain algorithm, and then counts the cells of each sample within each cluster. Louvain further uses the same procedure as Milo to ultimately determine the DA cells.

Due to the fact that the six DA testing methods employ diverse strategies to estimate DA cell populations, it is hard to find a single threshold applicable to all of the methods. To avoid such bias, we utilized the area under the receiver operator curve (AUC) score to objectively quantify the performance of the various DA testing methods. Noting that our datasets do not have ground truth DA labels, we employed a data-driven technique to construct such labels for each individual cell after setting the target DA cell populations. By aggregating results across all datasets with different experimental configurations, we ultimately ranked the overall predictive performance of the six DA testing methods. This gauges how well cell populations that are strongly related with the corresponding conditions, such as phenotype and experimental perturbations, can be inferred via DA testing.

In addition to accurately inferring the condition-specific cells, further issues should be addressed to make DA testing more applicable in real-world settings. First, in order to provide reliable predictions, a DA testing method must be robust to other variables in a dataset, such as batch effects and sample covariates. Second, a DA testing method should be resilient to datasets with different characteristics. For instance, (1) There are numerous single-cell profiling modalities, such as scRNA-seq and CyTOF to measure the expression of genes and proteins, respectively; (2) The structure of differential trajectories in single-cell datasets varies in high-dimensional gene or protein expression space; and (3) The size and differential abundance ratio of DA cell populations can vary between datasets. Finally, a practical DA testing method should be computationally efficient, such that it can be readily applied to single-cell datasets containing more than 100, 000 cells. Hence, we ran a series of experiments to evaluate and compare the performance of DA testing methods in various configurations and to see if they are capable of handling the challenges above. Furthermore, we also carried out studies to examine the sensitivity of results with respect to the input hyperparameters. This is crucial because, in practical applications, it might be difficult for users to specify an appropriate hyperparameter as input for a new given dataset without some background knowledge (Section Hyperparamter tuning and sensitivity). In our implementation, the hyperparameters were tuned as suggested in the original work. For specific hyperparameter values used in our experiments, refer to Supplementary Table S2.

### DA testing performance on synthetic datasets

First, the DA testing methods were evaluated on three synthetic datasets, referred to throughout the text as *linear, branch*, and *cluster* (see Supplementary Figure S1). Figure 3 and Supplementary Table S3 show the performance of the six DA testing methods on the three synthetic datasets and generally show how the performance changes with respect to DA ratios. On the synthetic datasets, we represented each individual high-dimensional cell in terms of its top 50 principle components (PCs), and systematically evaluated performance across varying DA ratios, RA target populations, and random seeds. When comparing the performance of the DA testing methods, we unsurprisingly observed that the accuracy of all DA methods as evaluated with AUC increased consistently as the DA ratio increased from 0.75 to 0.95 across the three datasets. This suggests that a higher DA ratio leads to a simpler and less-noisy DA testing problem (Figure 3). To quantify the performance of each method, we averaged the median AUC scores over all DA ratios. The average AUC scores of DA-seq, Meld, Milo, Cydar, Cna, and Louvain for the linear dataset were 0.91, 0.98, 0.98, 0.96, 0.79, and 0.80, respectively (Supplementary Table S3). On the linear dataset, the performance of the DA testing methods can be grouped into three major groups: (1) Meld, Milo, and Cydar; (2) DA-seq; and (3) Cna and Louvain. Meld and Milo were the most effective methods, and in all DA ratios, they performed slightly better than Cydar. DA-seq performed worse than the approaches in groups (1) but better than those in group (3). Similar patterns were also observed in the branch dataset (Figure 3B). The average AUC scores for Meld, Milo, and Cydar were 0.95, 0.95, and 0.93, respectively. The average AUC value for DA-seq was 0.90, which was lower than Meld, Milo, and Cydar but higher than the other approaches. The average AUC values of Cna and Louvain for the branch dataset were 0.76 and 0.79, respectively. Noting that the DA testing methods provided similar performance and relative performance rankings on the linear and branching datasets, we hypothesize that this was due to their similar differential trajectories.

**Figure 3:**
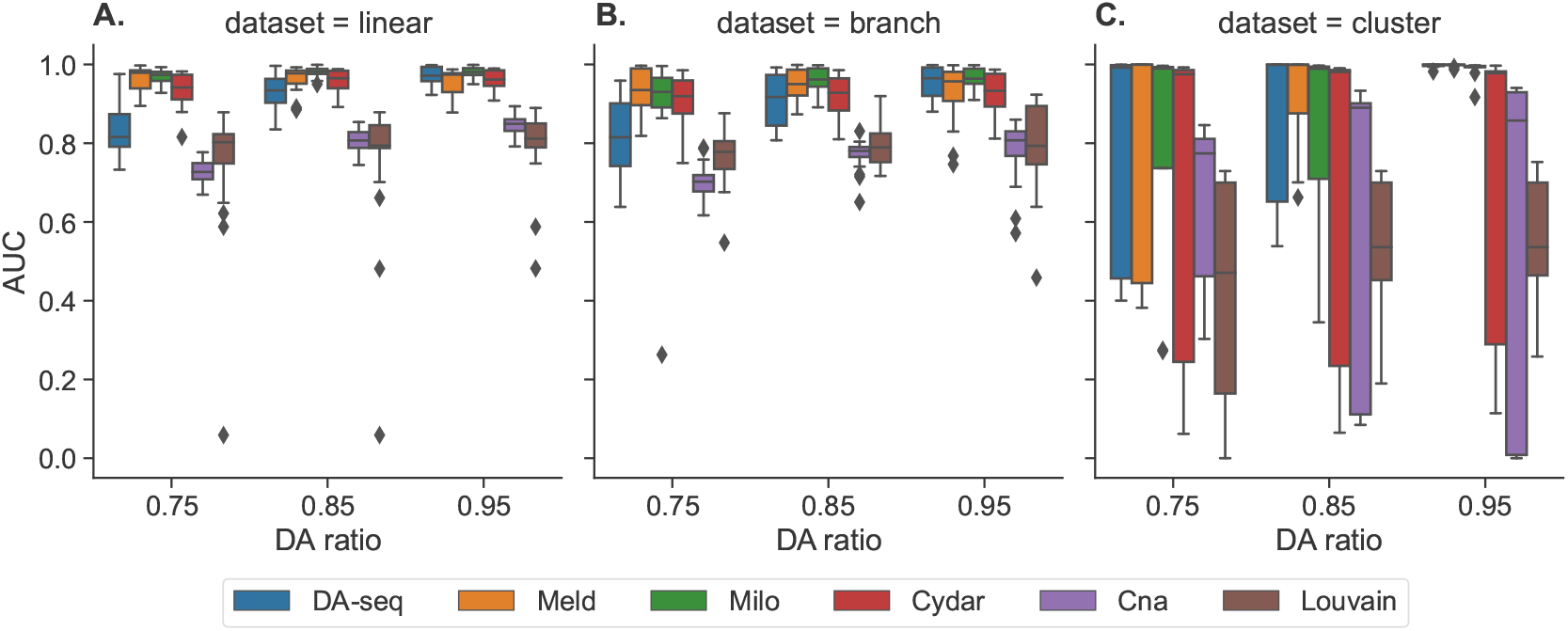
Performance of the six DA testing methods for DA prediction on the three synthetic datasets (linear (A), branch (B), and cluster (C)) with a range of DA ratios (0.75, 0.85, and 0.95) in the target DA cell population. The boxplots represent the distributions of AUC scores for various target DA cell populations across multiple random seeds.

In the cluster dataset, the averaged AUC scores of DA-seq, Meld, Milo, Cydar, Cna and Louvain were 1.00, 1.00, 0.99, 0.98, 0.84 and 0.51, respectively (Figure 3C). The average AUC values of DA-seq, Meld, Milo, and Cydar were higher than their results on the linear and branch datasets. When examining the distribution of AUC scores across various seeds, we found extremely high variance, especially for DA ratios of 0.75 or 0.85. In addition, the imbalanced distribution of AUCs (high median value and variance) in Figure 3C revealed that the DA testing methods performed very poorly in a small subset of the experiments. To further explore this, we visualized boxplots of AUC scores for each target DA cell population on the cluster dataset (Supplementary Figure S2). This suggested that the DA testing methods perform well and consistently when the target DA population is M1 cell type or M3 cell type, whereas for the M2 population (Supplementary Figure S2A), all the methods had a significant drop in performance compared to M1 and M3, indicating that the DA testing methods may not be effective in some special cases. In contrast to the linear and branch datasets, the cluster dataset has a more variable number of cells across populations. For example, the M2 population contained significantly more cells than the M1 and M3 populations. As a result, we hypothesized that the DA testing methods struggle when there is an imbalance and variable number of cells across populations, and specifically one such population contains substantially more cells than the others. To test this hypothesis, we subsampled the M2 population and ran a second experiment on a *balanced* cluster dataset. When the target DA population was M2 on the balanced cluster dataset, the performance of each method was greatly improved (Supplementary Figure S2B). In addition, compared to the results on the cluster dataset (Supplementary Figure S2A), the variance of AUC scores across the three target populations (M1, M2, and M3) was significantly reduced on the balanced cluster dataset, indicating that there were no strong biases between the various target DA populations after their numbers were balanced. Our experiments therefore revealed a possible limitation with respect to how the current DA testing methods are designed. Namely, these DA testing methods may not be able to adequately account for the biases caused by the imbalance in cell quantity across distinct cell populations and may therefore prioritize incorrect cells.

### DA testing performance on scRNA-seq and CyTOF Datasets

Next, we used an scRNA-seq and a CyTOF dataset to benchmark the DA testing methods. The first dataset is a scRNA-seq dataset termed COVID-19 PBMC [5], which was profiled from seven hospitalized patients at varying stages of COVID-19 development and another six healthy donors. The COVID-19 PBMC dataset consists of 44, 721 peripheral blood mononuclear cells (PBMCs) from 13 distinct cell types (Figure 4A) and the expression of 26, 361 genes. The second dataset is the BCR-XL mass cytometry dataset [8]. The BCR-XL dataset contains 172, 791 human PBMCs analyzed from 16 CyTOF samples, of which eight were stimulated with B cell receptor/Fc receptor cross-linker (BCR-XL). Originally, the BCR-XL dataset consisted of 35 different measured parameters. The cell types within the BCR-XL dataset were manually gated using some predefined phenotypic markers (Figure 1A). Note that these two datasets come from two distinct modalities. The COVID-19 PBMC dataset profiles the transcriptome, whereas the BCR-XL is a single-cell proteomics dataset. In addition, scRNA-seq datasets contain significantly more features than cytometry datasets (e.g., 26, 361 vs. 35). Consequently, the datasets we chose can adequately represent standard and widely used single-cell datasets.

**Figure 4:**
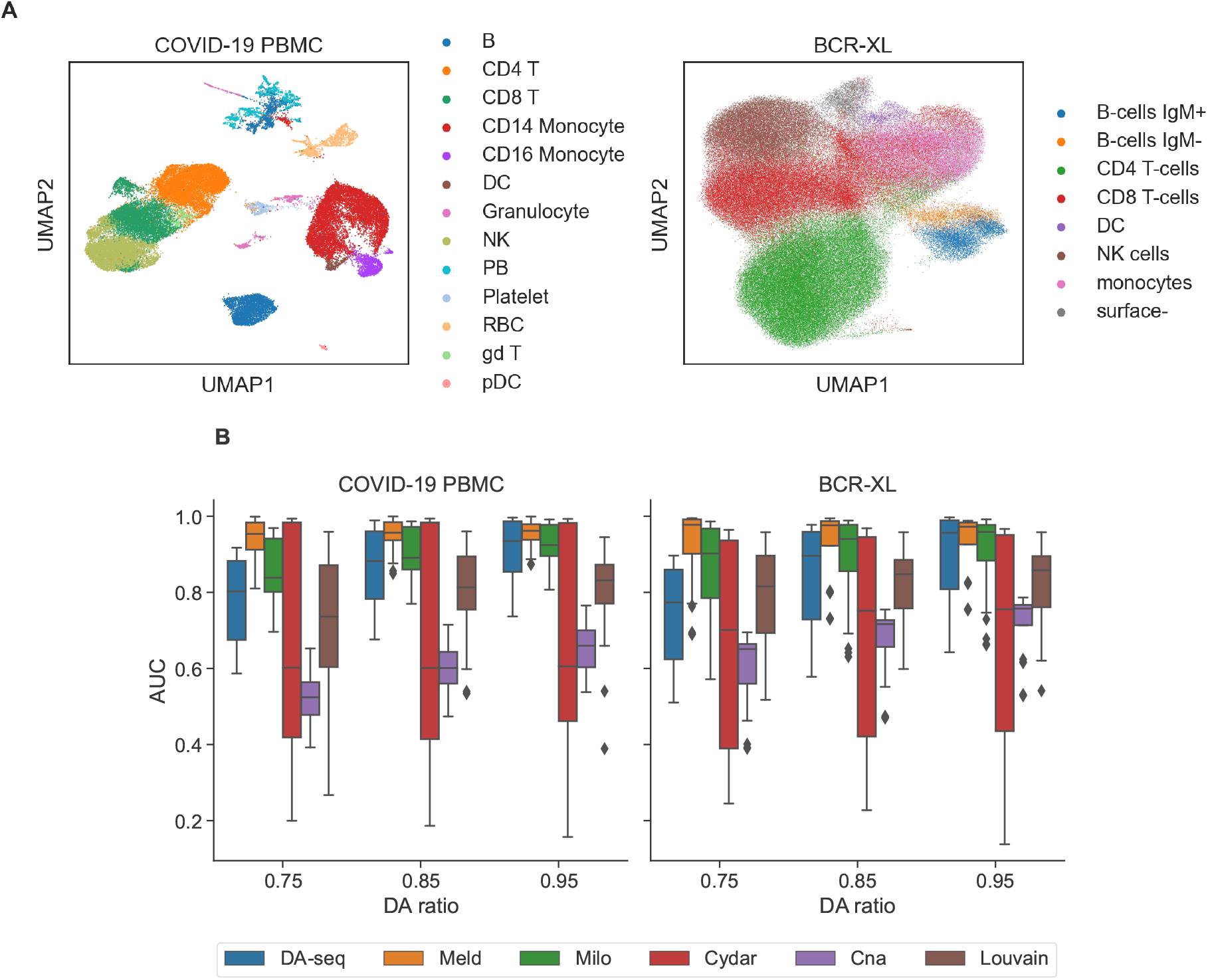
(A) UMAP visualization of the cells in the COVID-19 PBMC scRNA-seq (left) and BCR-XL CyTOF datasets (right), colored according to annotated cell-types. (B) Performance of the six DA testing methods for DA prediction on the two real single-cell datasets (COVID-19 PBMC (left) and BCR-XL (right)) with a range of DA ratios (0.75, 0.85, and 0.95) in the target DA cell type. The boxplots represent the AUC scores for different target DA cell types (in A) evaluated over different random seeds.

In the tests with the COVID-19 PBMC and BCR-XL datasets, we applied the same evaluation procedure as in the synthetic datasets. That is, we selected each cell type as a target DA cell population and evaluated the quality of each of the six DA testing methods across various DA ratios and three random seeds. We also generated the ground truth DA labels for each cell in the two real datasets similarly to how we did with the synthetic datasets. To reduce the computational complexity of the COVID-19 PBMC dataset, we used the top 50 PCs as input. For the BCR-XL dataset, we used its filtered raw features as input. Figure 4B and Supplemental Table S4 show the benchmarking results on the COVID-19 PBMC and BCR-XL datasets. Consistent with the pattern of the synthetic datasets, we also observed that the performance of all methods improved steadily as the DA ratio increased. Similarly, the variance of AUC scores also decreased as the DA ratio increased across datasets for all methods except for Cydar. This showed that, similar to the patterns observed in the synthetic dataset, the DA ratio can significantly affect the performance and stability of the DA testing methods. In the COVID-19 PBMC dataset, the mean AUC values for DA-seq, Meld, Milo, Cydar, Cna, and Louvain were 0.87, 0.96, 0.88, 0.60, 0.59, and 0.79, respectively. Meld ranked first among the six DA testing methods applied to the COVID-19 PBMC dataset, followed by Milo, DA-seq, Louvain, Cydar, and Cna, with Milo and DA-seq performing similarly, and Cydar and Cna also performing similarly. In the BCR-XL dataset, the corresponding average AUC values for DA-seq, Meld, Milo, Cydar, Cna, and Louvain were 0.88, 0.98, 0.93, 0.74, 0.71, and 0.84. As a result, Meld was ranked first among the six DA testing techniques on the BCR-XL dataset as well, surpassing Milo, DA-seq, Louvain, Cydar, and Cna. Overall, the methods’ performances and rankings remained consistent in both the synthetic and real single-cell datasets, demonstrating that their performances are independent to the data but are reflections of their own capabilities. In addition, the performance of DA-seq, Meld, Milo, and Louvain on the COVID-19 PBMC and BCR-XL datasets was comparable to their performance on synthetic datasets, but Cydar and Cna showed a considerable fall in performance. This demonstrated that (1) DA-seq, Meld, Milo, and Louvain were more adaptable to different kinds of data, like synthetic, scRNA-seq, and CyTOF single-cell datasets; and (2) Cydar and Cna may not have been as adept at adjusting to the biases between synthetic and real datasets, or they may have been sensitive to changes in other factors, such as hyperparameters.

### DA testing performance on datasets with additional technical and biological covariates

In addition to the clinical outcomes used in DA testing, such as clinical phenotype or disease status, single-cell datasets are often affected by additional technical and biological factors, such as batch effects, donor type, and cell cycle artifacts. These undesirable variables provide additional variance in the data and can confound the biological variations of relevance in the subsequent analysis, resulting in more false positives. In this subsection, we examine how the performance of DA testing methods changes when batch effects are present in the data, as well as how each DA testing method particularly accounts for additional covariates, such as batch effects. Out of the six DA testing methods, Cydar, Milo, Cna, and Louvain can explicitly include such external variables into their testing models to account for variance, whereas DA-seq and Meld do not. Following the procedure in Ref. [14], we simulated batch effects with a variety of magnitudes and added them to the three synthetic datasets. We then evaluated the performance of the six DA testing methods on the synthetic datasets containing batch effects (Figure 5). To exclude influence from different hyperparameters, we used the same hyperparameter values as when no batch effects were present. Overall, the results demonstrated that all DA methods exhibited poorer performance when batch effects were present in the data, in comparison to batch-effect free data, as exhibited by a decline in AUC scores with increasing magnitude of batch effects (Figure 5). This showed that technical artifacts such as batch effects had a significant adverse effect on the quality of DA testing methods. Of the six DA testing methods, Milo consistently performed the best over a range of batch effect magnitudes. Despite no explicit implementation to include batch labels in their models, DA-seq and Meld were inferior to the performance of Milo but outperformed the other approaches. We also noticed a strong negative link between how well DA-seq and Meld worked and how strong the batch effects were (Figure 5). Cydar, Cna, and Louvain were the weakest methods for handling batch effects as their performances were affected by even slight batch effects. Given that Cydar, Cna, and Louvain models covariate in the same way as Milo, their low performance was more likely due to their method-specific cell counting step. In addition, we conducted a second experiment to examine whether incorporating batch labels into the models could improve the performance. In this experiment, we applied Cydar, Milo, Cna, and Louvain on synthetic datasets using two different setups. In the first setup, batch labels were included in the models, whereas in the second setup, they were omitted. Supplementary Figure S3 illustrates the performance of these four approaches with or without inclusion of batch information. We discovered that, with the exception of Cydar, explicitly modeling batch effects can greatly enhance the performance of DA testing procedures in datasets with prominent batch effects. Furthermore, these experiments demonstrated that it is crucial for DA testing procedures to account for the variance introduced by additional technical and biological factors in order to produce accurate and meaningful results.

**Figure 5:**
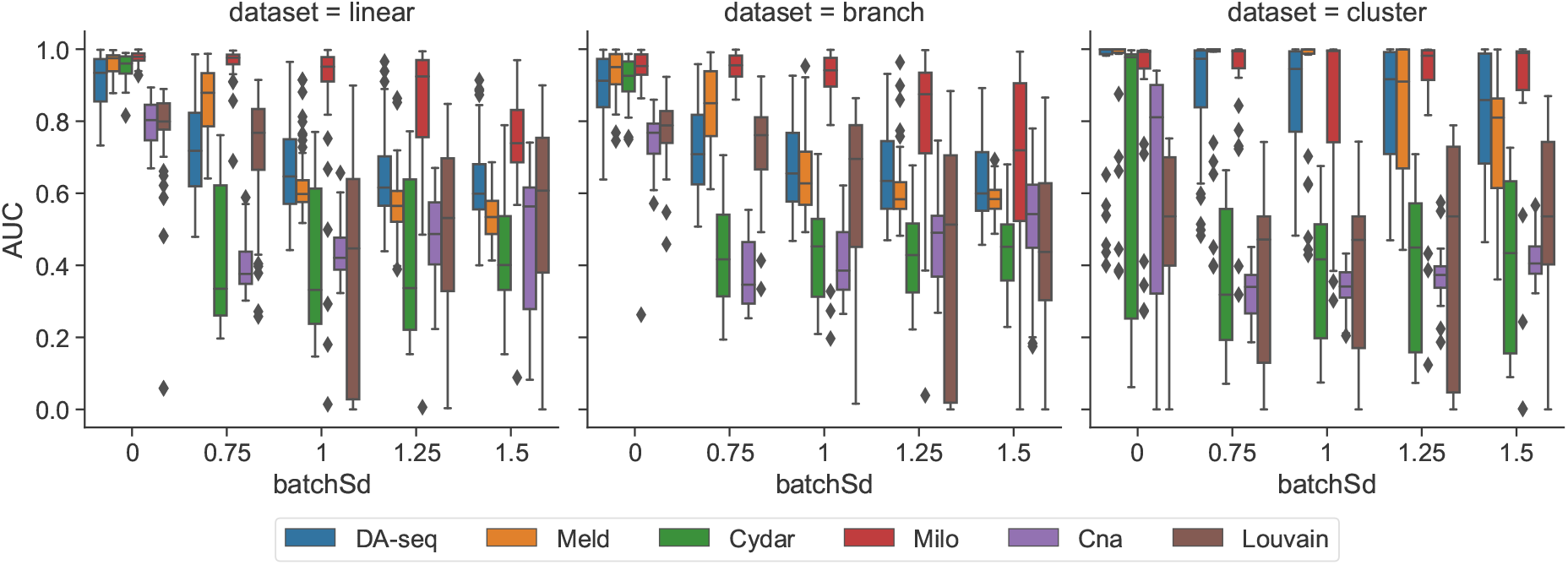
Performance of the six DA testing methods for DA prediction on the three synthetic datasets (linear (left), branch (middle), and cluster (right)) with batch effects of varying magnitudes (from 0 to 1.5). When `batchSd=0`, no batch effects are present. The boxplots represent the AUC scores for different target DA cell populations, DA ratios, and random seeds.

### Runtime efficiency and scalability of DA testing methods

Next, we evaluated the runtime efficiency and scalability of the six DA testing methods. We measured the execution time of each method on the COVID-19 PBMC and BCR-XL datasets using the tuned hyperparameters. In the BCR-XL dataset containing more than 170,000 cells, all methods were able to finish running within a few hours (Figure 6A). Additionally, some of the DA testing approaches with even higher efficiency, such as Cydar, Cna, and Louvain, only took a few minutes. Thus, we proved that efficiency is not a limiting factor for any of the six DA testing methods when applied to the vast majority of single-cell datasets. Noting that wall-clock runtime depends on numerous factors such as algorithm complexity, hyperparameters, and computing infrastructure, it cannot objectively and completely reflect the scalability of the DA testing methods. As a result, we conducted an additional experiment to quantify the scalability of the DA testing techniques by evaluating the relative runtime growth rate as the number of cells increased. Using the R package splatter [22], we constructed six single-cell datasets with increasing numbers of cells (4k, 10k, 15k, 30k, 50k, and 100k). To specifically evaluate the runtimes of the core components of the DA methods without the variable times required for hyperparameter selection, we used default hyperparameters for all datasets. We quantified how well each method scaled by calculating the relative runtime growth with respect to the runtime on the smallest dataset with 4k cells for various data sizes. Figure 6B shows the relative runtime growth as a function of the number of cells in each dataset. Among the six DA testing methods, we discovered that Cna was the most scalable, while Louvain and Meld were the least scalable. The scalability of DA-seq, Milo, and Cydar fell between Cna and Meld, with Cydar being marginally superior to Milo and DA-seq. Notably, the majority of the runtime of the six approaches was spent either counting cells across conditions in cell neighborhoods or building the cell-to-cell graph across samples.

**Figure 6:**
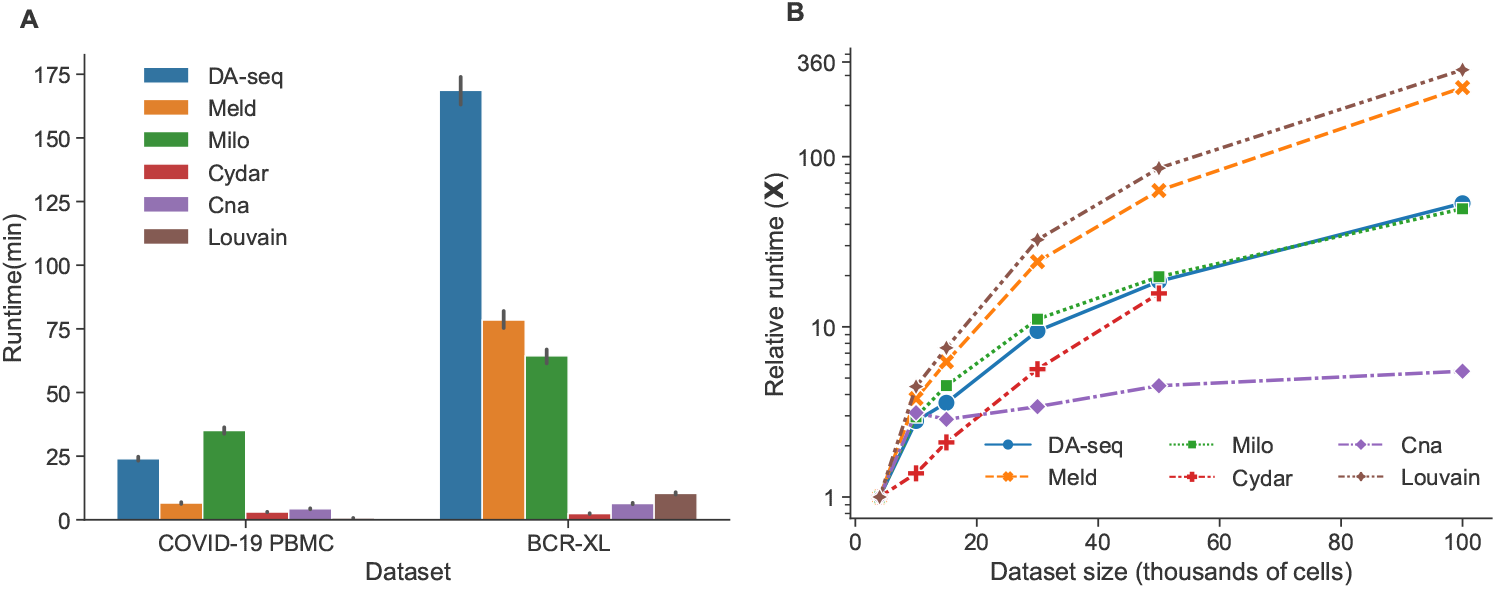
Runtime efficiency and scalability of the six DA testing methods. (A) Runtime of the six DA testing methods on the COVID-19 PBMC (left) and BCR-XL (right) single-cell datasets. The error bars reflect the standard deviation of runtime across various target DA cell populations, DA ratios, and random seeds. The execution times were measured on the nodes of a cluster with Intel Xeon E5-2680 v3 CPUs and 256GB RAM. (B) Relative runtime growth ratio of the six DA testing methods on the single-cell datasets as a function of an increasing number of cells (4k, 10k, 15k, 30k, 50k, and 100k). The runtime of the smallest dataset was used to normalize the runtimes of the larger datasets.

### Hyperparamter tuning and sensitivity

We further evaluated the hyperparameter tunability and sensitivity of the six DA testing methods. Hyperparameters are crucial to the performance of machine learning methods and the best way to identify the optimal hyperparameters is through a line or grid search in hyperparameter space, which takes a lot of time and computational resources. In general, machine learning models with fewer hyperparameters are easier to tune. Furthermore, if a machine learning model’s performance is sensitive to its hyperparameters, it is challenging to identify the best hyperparameters, hence making the model’s performance unstable. Thus, we propose to examine two criteria to evaluate the DA testing methods, including the number of hyperparmaters and the overall sensitivity of hyperparameters. The number of hyperparameters reflects how easily a method can be tuned, and sensitivity of hyperparameters measures how stable the DA testing method is overall. In Supplementary Table S5, we outlined the hyperparameters of the six DA testing methods. Milo, Cydar and Cna only have one hyperparameter, *k*, which is the number of *k*-nearest neighbors to use in the graph-representation of the data. Alternatively, DA-seq, Meld, and Louvain have more hyperparameters (Supplementary Table S5). As a result, the hyperparameters of Milo, Cydar, and Cna are easier to tune than those of DA-seq, Meld, and Louvain.

To test the hyperparameter sensitivity of each DA testing method, we evaluated their performance for predicting DA cells on three synthetic datasets (linear, branch, and cluster) and on the COVID-19 PBMC scRNA-seq dataset by altering their hyperparameters. Since Meld, Milo, Cna, and Louvain all share a common hyperparameter, *k*, we fixed *k* and solely tested the hyperparameter sensitivity relative to the other parameters in order to control variable. In contrast to other methods, DA-seq employs a range of hyperparameters **k** = [*k*_1_,…, *k*_l_] to generate *k*-nearest neighbor graphs. We altered the hyperparameters of DA-seq by replacing *k*_1_ with the same *k* used in the other methods, while varying the step size between *k_i_* and *k*_*i*+1_. The boxplots in Figure 7 visualize the variation in performance of DA-seq, Meld, Cydar, and Louvain, with each dot representing a run with specific hyperparameters. First, DA-seq had the lowest overall hyperparameter sensitivity, indicating that users do not need to modify its hyperparameters excessively for practical applications. Second, although having somewhat higher variance than DA-seq, Meld’s performance did not show a strong variance. Cydar and Louvain, on the contrary, consistently had high variance in their performances across all datasets. This experiment proved that Cydar and Louvain are hyperparameter-sensitive. The rationale for Cydar’s high hyperparameter sensitivity is because finding an appropriate radius in high-dimensional space is inherently difficult, as data points become sparser as the dimensionality increases [24]. Louvain’s strong hyperparameter sensitivity is primarily due to the resolution parameter, which ultimately controls the number of clusters identified. Taken together, it is crucial to identify the optimal hyperparameters for Cydar and Louvain for real-world applications; otherwise, these algorithms may perform poorly in certain circumstances.

**Figure 7:**
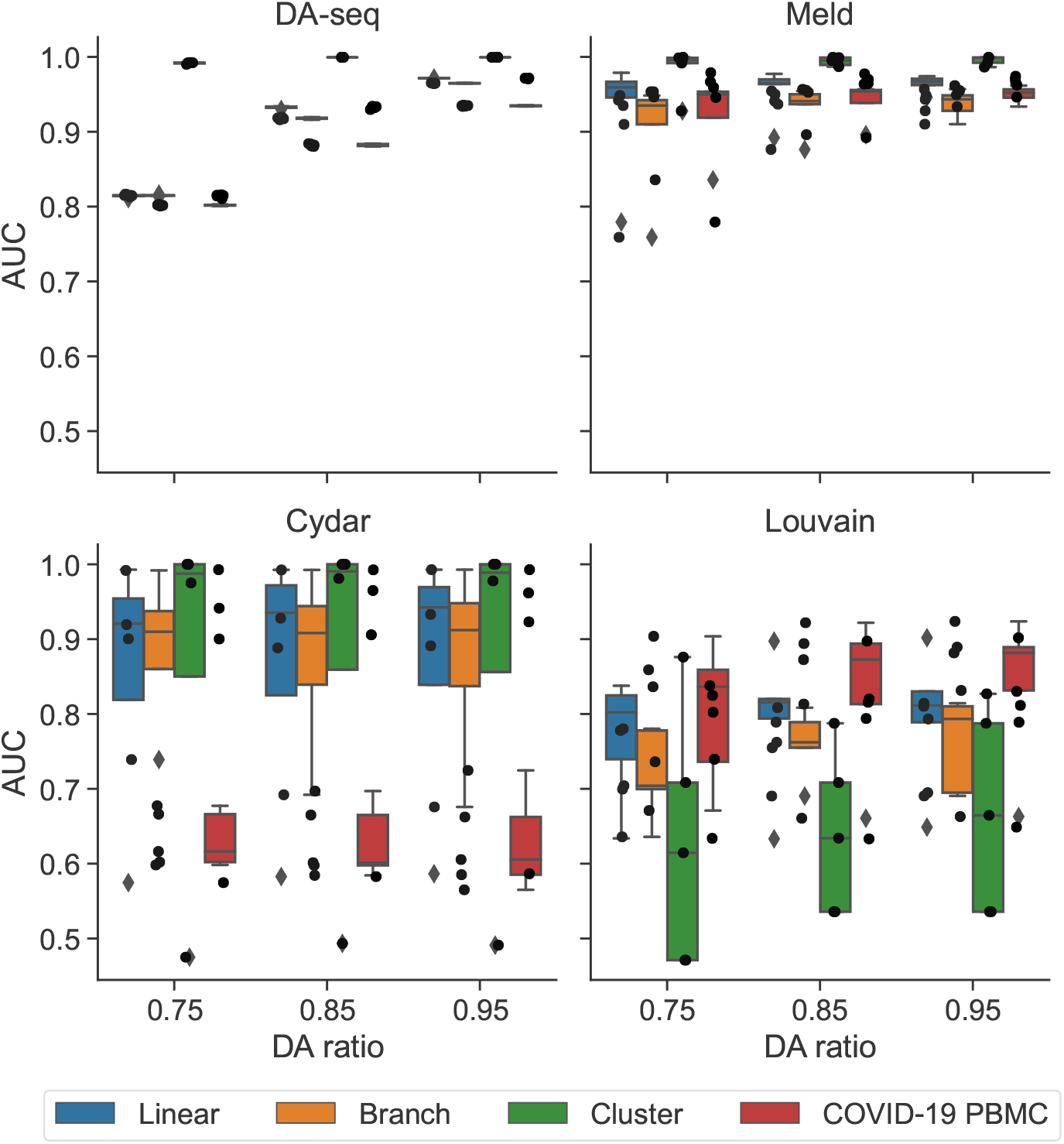
Hyperparamter sensitivity of the six DA testing methods. Performance of the four DA testing methods (DA-seq, Meld, Cydar, and Louvain) on the three synthetic datasets and the COVID-19 scRNA-seq dataset with a range of DA ratios (0.75, 0.85, and 0.95) in the target DA cell population. The boxplots show (e.g. each data point) the distribution of AUC scores across various hyperparameters. High variance implies sensitivity to choice of hyperparameters.

## Discussion

In this work, we evaluated and compared six prominent DA testing methods for resolving cell populations in response to external variables, such as clinical phenotype or experimental perturbation. Our benchmarking workflow was designed to cover as many realistic applications of DA testing scenarios as possible, including diverse single-cell data types, various data topologies, and the existence of technique-induced biases. In our experiments, we assessed the DA testing methods using both synthetic and real single-cell datasets with distinct topological structures. In addition, simulated batch effects were generated and applied to the datasets to assess the robustness of DA testing methodologies. Thus, our benchmarking strategies offered a thorough, quantitative evaluation of the DA testing methods. We evaluated the performance of each method by calculating AUC scores to quantify the similarity between predicted and established ground-truth DA labels. By objectively comparing the performance of the six DA testing approaches on a variety of tasks, we determined that no single method outperformed the others across the board. In other words, the appropriate selection of DA testing methods depends on properties of the data and ultimate task of interest (e.g. the existence of batch effects). In the discussion that follows, we summarize our experimental findings for each given task.

The majority of the DA testing methods examined in our work, particularly Meld, Milo, and Cydar, demonstrated consistently strong accuracy across datasets and DA ratios for identifying DA cell types in synthetic datasets (Figure 3 and Supplementary Table S3). Meld performed the best across all approaches for the real-world single-cell datasets, but Milo and DA-seq also attained satisfactory accuracy. When additional technical challenges, such as batch effects were present in the datasets, Milo was the most effective at correcting them and reducing their negative impacts on testing accuracy. In addition, we demonstrated that including batch labels in a DA testing model enhanced performance in comparison to not including them. As for runtime efficiency and overall scalability, all methods can successfully complete their workflows on typically-sized scRNA-seq and large CyTOF datasets using standard CPUs in a few hours. Finally, we looked at the sensitivity and usability of the hyperparameters used in the DA testing methods. While Milo required tuning of the fewest hyperparameters, DA-seq and Meld were robust to the selection of hyperparameters. Furthermore, our benchmarking evaluations identified a common problem across the majority of DA testing methods. In particular, all methods performed poorly, even on simple datasets, when a substantial imbalance of cells existed between cell-types. Our hypothesis is that such behavior is caused by the several data scaling options, which are intended to normalize the data. As this issue is seemingly complex, we leave a more in-depth analysis of this phenomenon across methods and datasets to our future work.

Based on our thorough benchmarking analyses, the following are our general suggestions for the usage of DA testing methods in practice (Supplementary Table S6). First, we observed that Meld is the overall most accurate method when there is no substantial technical noise, such as batch effects. Moreover, in the event of technical or biological noise, Milo performs better on average than Meld. Our experiments further suggested that Milo, Cydar, Cna, and Louvain are all viable candidates for robustly identifying DA cell-populations, while controlling the false discovery rate. Milo, DA-seq, and Meld either have the fewest hyperparameters or are insensitive to hyperparameter changes. Therefore, they are the robust strategies for performing DA testing on a new dataset. Lastly, for large single-cell datasets with many cells, we found Cna to be the most scalable method; if long run times in your analysis are intolerable, we advise you to attempt scalable DA testing methods, such as Cna. We hope that the presented benchmarking study will assist users in selecting the optimal method for their DA testing tasks.

## Methods

### Data Pre-processing

For all the synthetic datasets, the raw count matrices created by the `generate_dataset()` function in dyntoy [21] were first normalized using a `log+1` transformation. Then, we projected the normalized gene expression data into principal component (PC) space and embedded the data into the manifold approximation and projection (UMAP) [25] space using the pre-computed top-50 PCs. In our analysis of the COVID-19 PBMC dataset, we utilized the previously processed data introduced in Ref. [5], whose processing procedures adhered closely to best practices for the analysis of single-cell data [26]. In addition to the data, the authors also provided embeddings for each cell according to both PCA and UMAP. The PC embeddings were generated using the normalized data of the highly variable genes and the UMAP embeddings were constructed using the top 50 PCs. In the BCR-XL CyTOF dataset, we eliminated 11 nonfunctional markers and used the 24 remaining functional markers (CD3, CD45, pNFkB, pp38, CD4, CD20, CD33, pStat5, CD123, pAkt, pStat1, pSHP2, pZap70, pStat3, CD14, pSlp76, pBtk, pPlcg2, pErk, pLat, IgM, pS6, HLA-DR, CD7) in our experiments. The markers in the BCR-XL dataset were normalized using an arcsinh transformation with a cofactor of 5, as suggested in [27].

#### Differential abundance (DA) testing methods

##### Problem Formulation

We define a sample **X**_*n*_×_*p*_ = (**x**_1_,…, **x**_*n*_)^⊤^ to be the normalized gene or protein expression matrix with *n* cells and *p* measured features, where **x**_*i*_ = (*x*_1*i*_,…,*x*_*pi*_)^⊤^ is the feature vector for cell *i*. Given a collection of *N* samples 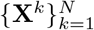 profiled from *N* individuals (donors), with a particular sample *i* associated with a label *y_i_*, the goal of DA testing is to identify a subset of cells exhibiting differential abundance (density) in response to the labels encoded (e.g. *y_i_* for individual *i*) across the samples. This DA testing problem can alternatively be stated as a density estimation problem [24]. In this case, each experimental condition can be viewed as a primary distribution, and the objective of DA testing is to detect cells with relatively lower or higher densities under each label or condition. In this subsection, we describe all of the benchmarking methods in this study. For a more detailed introduction about the DA testing methods, please refer to their respective original papers [23, 13, 16, 10, 14, 15].

##### Cydar

Cydar [13] is a statistical testing approach developed to identify cell populations in single-cell mass cytometry data with a differential abundance of cells between conditions. Cydar’s central idea is to construct hyperspheres in the multi-dimensional marker space as local units to test if the number of cells among samples in each hypersphere is related to external labels, such as clinical or experimental outcomes. Given an *N*-sample single-cell dataset measuring *p* markers in each cell, Cydar’s testing pipeline works as follows: (1) Cydar randomly samples a subset of cells from the entire dataset and uses these cells as the centers of hyperspheres to allocate cells from all samples to the hyperspheres; (2) Cydar then counts the number of cells assigned to each hypersphere in each sample, resulting in an *N*-dimensional abundance vector; (3) Next, Cydar employs the negative binomial generalized linear models (NB-GLMs) in edgeR [12] to perform statistical testing on these count data with respect to clinical outcomes and other informational covariates, and assigns a *P*-value to each hypersphere; (4) Lastly, Cydar identifies the statistically significant hyperspheres as DA regions by controlling the spatial false discovery rate (FDR), a weighted form of FDR that regulates FDR across volume, at a predetermined threshold *α*. Here, Cydar applies the Benjamini-Hochberg (B-H) procedure [28] to calculate the maximum P-value needed to keep a hypersphere below the spatial FDR threshold *α*, which is defined as,

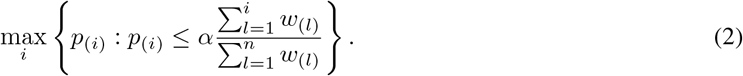

Here, *n* is the number of hyperspheres, *p*_(1)_ < *p*_(2)_ <… < *p*_(*n*)_ order the *P*-values of the hyperspheres and *w*_(*l*)_ is defined as weight, which is the reciprocal of the density of hypersphere (*l*). In our benchmark, Cydar v1.18 (http://bioconductor.org/packages/cydar) was applied across all the experiments.

##### DA-seq

In DA-seq [16], a logistic regression classifier is used to compute a local DA score for each cell so that DA subpopulations can be identified. The logistic regression classifier takes the cells’ feature vectors as input, which measure the abundance of two biological conditions in the area around each cell at different scales. DA-seq trains the logistic regression classifier by using cells’ condition labels and the feature vectors. The fitted probability is then used as the DA score for each cell. In this case, the trained logistic regression model serves as a smoothing function that transforms a cell’s input feature vector to its corresponding soft DA score. Next, DA-seq uses a random permutation test to find statistically significant DA cells in the dataset. The upper and lower cut-off thresholds are based on the highest and lowest DA scores inferred under the null hypothesis that the condition labels are distributed randomly. In our experiments, we used the official DA-seq implementation, which can be accessed at https://github.com/KlugerLab/DAseq.

##### Meld

Meld [10] is a graph-based kernel density estimation method. It is used to estimate the likelihood of a sample (often referred as a cell) under various experimental perturbations. Inspired by the recent success of applying manifold learning techniques to single-cell data visualization [25, 29, 30], Meld extends kernel density estimation (KDE) from the regular spatial domain to a manifold represented by a cell-by-cell similarity graph denoted by 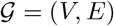. Here, Meld requires two steps to obtain the edge weights in 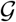. First, the Euclidean distance between cells is calculated for a pair of cells, (*i*,*j*). Next, the weight (similarity) between a cell pair (*i*,*j*) by feeding their distance to some predefined kernel functions, such as the *α*-decaying kernel [30] or the MNN kernel [31].

The Meld algorithm interprets the cell label as a signal across the cell-cell similarity network. It employs a low-pass graph filter [32] to de-noise the node labels across the graph and uses the smoothed label as the DA score measurement for each cell. Noting that this graph filtering step is performed independently on each condition, the smoothed condition labels for each cell must be normalized (summed to 1) in order to derive the conditional label associated likelihood. For experiments with several experimental and control duplicates, the Meld algorithm must be applied to each replicate separately, and the DA scores therefore must be averaged across replicates. Meld uses a heuristic strategy to choose DA cell subpopulations by setting a threshold on the per-cell likelihoods to determine whether a cell is in a zone where a certain label is more or less abundant. We used the Meld python package, which can be accessed at https://github.com/KrishnaswamyLab/MELD.

##### Cna

Cna, or “co-varying neighborhood analysis”, identifies phenotype-associated cell populations by examining cell neighborhoods that co-vary in abundance with respect to certain sample covariates, such as experimental treatment or clinical outcome. Similar to Meld, the Cna approach begins by constructing a *k*-nearest neighbor graph of cells across all samples. Cna adopts the `scanpy.pp.neighborhood()` function from the `scanpy` package to encode the neighborhood associations between cells into a sparse weighted adjacency matrix **A**. Next, Cna uses a random walk to calculate the likelihood that the *m*′-th cell is in the neighborhood of the *m*-th cell. Formally, this is given by

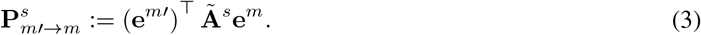

Here, s represents the steps of random walk, **e**^*m*^ and **e**^*m*′^ are the indicator vector defined at indices *m* and *m′*, respectively, and **Ã** is the random-walk markov matrix with self-loops, whose entries are computed as,

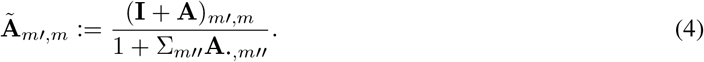

Here, **I** is an identity matrix and **A** is the weighted adjacency matrix that is computed in the graph building step. Letting *c*(*n*) denote the cells from sample *n*, then **R**_*n*,*m*_ is the expected number of cells that would arrive at the neighborhood of the *m*-th cell after *s* steps of random walking beginning from sample *n*. Formally this is calculated via 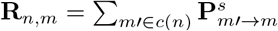. Cna further defines the neighborhood abundance matrix (NAM) **Q** ∈ ℝ^*n*×*m*^ by normalizing the rows of **R** (summed to 1), where

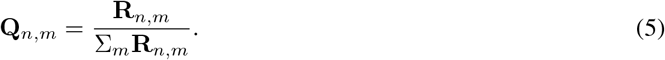

Once the NAM is defined, Cna tests its association with a known sample-level covariate **y** using a linear regression model. The linear model is formally defined as,

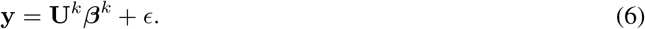

Here, **U**^*k*^ represents the first *k* columns of **Q**’s left matrix of singular vectors **U**, *β^k^* is the vector of coefficients, and *ϵ* denotes zero-mean Gaussian noise. Thus, the *P*-value is calculated using a multivariate *F*-test for a range of *k*s, such that the one attaining the smallest *P*-value is ultimately selected. To identify the differentially abundant neighborhoods, Cna computes a “smoothed correlation” between each neighborhood *m* and the sample-level covariate **y**. The smoothed correlation is mathematically defined as,

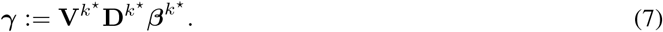

Here, *k*^*^ denotes the optimal number of singular vectors (e.g. components) determined by the multivariate *F*-test, 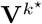 is the first *k*^*^ columns of **Q**’s right singular vector matrix, 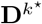 is the top-left *k*^*^ × *k*^*^ submatrix of **Q**’s singular vector matrix and 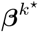 is the coefficient vector defined in 6. To assess the statistical significance, the null distribution of *γ* is obtained by fitting 6 using different permutations of **y**. Lastly, the DA cell sub-populations are determined by a given FDR threshold for *γ*. The Cna approach is implemented in python, and is available at https://github.com/immunogenomics/cna.

##### Milo

As an improved version of Cydar, Milo also uses NB-GLMs to test DA cells in single-cell datasets but replaces the hypersphere in Cydar with cell neighborhoods from the cell-cell similarity graph. Here, the neighborhood of a cell *c_i_* is defined as the set of first order neighbors(including *c_i_* itself) in a *k*NN graph created by the `findKNN()` function in the `BiocNeighbors` package. After counting the number of cells in the neighborhoods of several samples, Milo employs the same statistical testing pipeline with `edgeR` as Cydar, except that the testing unit is a cell-neighborood instead of a hypersphere. To reduce complexity, Milo samples only a small proportion (by default, 0.1) of cell neighbors to find DA neighborhoods. As various cell neighborhoods may share certain cells in the *k*NN graph, it is vital to highlight that a cell neighborhood must propagate its DA score to each of its respective cells. Hence, the DA score of a tested cell is ultimately determined by adding the DA scores of all the cell neighborhoods to which it belongs. In this work, we used the `R`-based implementation of Milo (https://github.com/MarioniLab/miloR), as suggested by the authors.

##### Louvain

The Louvain algorithm [23] is a cluster-based approach for DA testing. Unlike other more granular approaches, which are performed on single cells [16, 10], hyperspheres [13], and cell neighborhoods [14, 15], Louvain’s results are typically coarser, operating on a cluster level, and hence can only determine whether a cell cluster is a DA region or not. In other words, if a cell cluster is determined to be a DA cluster, all cells insides this cluster become classifed as DA cells with identical DA scores. The Louvain method is implemented as follows: (1) a *k*NN graph or cell-to-cell similarity graph is constructed; (2) the Louvain algorithm partitions the graph into clusters [23] (implementation provided by the `cluster_louvain()` function in the `R` package `igraph` [33]); and (3) apply the statistical framework of Milo [14] to identify DA cell-populations. Note that the Louvain approach does not implement DA-score aggregation step introduced by Milo and therefore produces solely non-overlapping cell clusters.

##### Evaluation and Metrics

Evaluating and comparing the performance of the various DA testing methods is non-trivial due to their variable testing procedures for identifying and quantifying the significance of DA cells. Cydar, Milo, and Cna, for instance, employ traditional statistical testing measures like FDR and spatial FDR to detect DA cell populations, whereas DA-seq and Meld use a conditional probability threshold. Therefore, it is impossible to develop a uniform criterion that can be consistently applied to all methods. To eliminate the bias of selecting a distinct threshold for each approach, each method’s predicted labels were generated using a range of thresholds based on its own criterion. To quantify overall classification performance, we compared predicted labels with ground truth labels that were generated through simulation for each cell and had three distinct categories: (1) enriched in C1 (NegLFC, negative log fold-change in condition C2 vs. C1), (2) enriched in C2 (PosLFC, positive log fold-change in condition C2 v.s. condition C1), and (3) Not DA, respectively.

To generate a list of evaluation thresholds, we first calculated DA scores (FDR or conditional probability, depending on the approach) under each method. Next, for each method, we specified the thresholds using the values at different percentiles (by default: 0% to 100% with 1% increments) of its DA scores. We then used the false positive rate (FPR) and true positive rate (TPR), two binary classification metrics, to assess the performance of each approach for each threshold, yielding a list of FPR and TPR pairs. We treated both the PosLFC and NegLFC as the “positive” label of binary classification to account for the fact that there are three possible ground-truth labels. The FPR and TPR are defined respectively as,

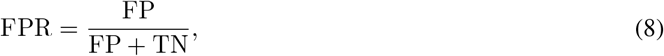

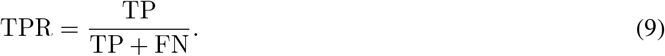

Here, FP is the number of cells with false positive DA predictions, TN is the number of cells with true negative predictions, TP is the the number of cells with true positive predictions, and FN is the number of cells with false negative predictions. Finally, we connected the 2d-points of the FPR and TPR pairs sequentially with FPR plotted on the horizontal axis and TPR plotted on the vertical axis to construct receiver operator curves (ROC) and reported the area under ROC (AUC) score as the overall performance for each method.

## Data and code availability

The raw data for the COVID-19 PBMC single-cell RNA-sequencing dataset is publicly available through the NCBI Gene Expression Omnibus with the accession number GSE150728. The authors of the original work also provide processed count matrices with manually annotated metadata and pre-computed embeddings in .`h5ad` and .`rds` formats, which can be downloaded from the Wellcome Sanger Institute’s COVID-19 Cell Atlas at https://www.covid19cellatlas.org/#wilk20. The raw data of the BCR-XL dataset including FCS files and their corresponding metadata and annotations can be downloaded from FlowRepository with experimental id: FR-FCM-ZYL8. All code to reproducethe results are available at https://github.com/CompCy-lab/benchmarkDA.

## Authors’ contributions

HD, NS, AP conceptualized and designed the study. HD performed data preprocessing, benchmarking, evaluation, and analysis. HD wrote the manuscript with input from all authors. All authors read and approved of the final manuscript.

## Acknowledgements

The authors thank the anonymous reviewers for their valuable suggestions.

