## Supplemental figures and tables for "Benchmarking differential abundance methods for finding condition-specific prototypical cells in multi-sample single-cell datasets"

### Benchmarking differential abundance methods for finding condition-specific prototypical cells in multi-sample single-cell datasets: supplementary figures and tables

HAIDONG YI, ALEC PLOTKIN, NATALIE STANLEY

#### 1. SUPPLEMENTARY FIGURES

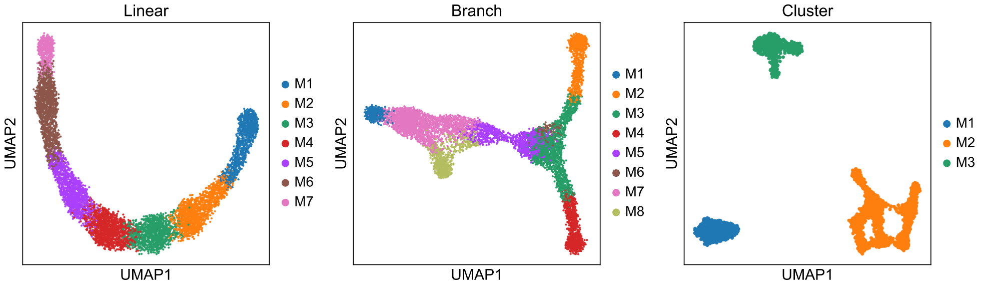


**Figure S1.** Two-dimensional UMAP visualization of the three synthetic single-cell datasets, where the cells are colored by target DA cell populations (cell type).

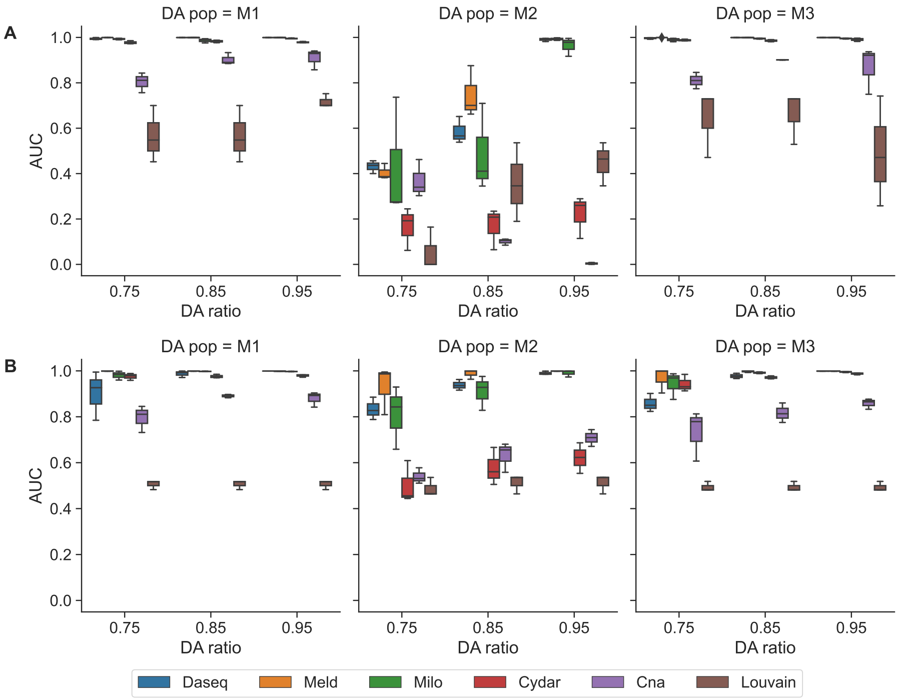


**Figure S2.** Performance of the six DA testing methods for DA prediction on the cluster dataset (A) and the *balanced* cluster dataset (B) with a range of DA ratios (0.75, 0.85, and 0.95) in the target DA cell populations (M1, M2, and M3). The boxplots represent the AUC scores for different random seeds.

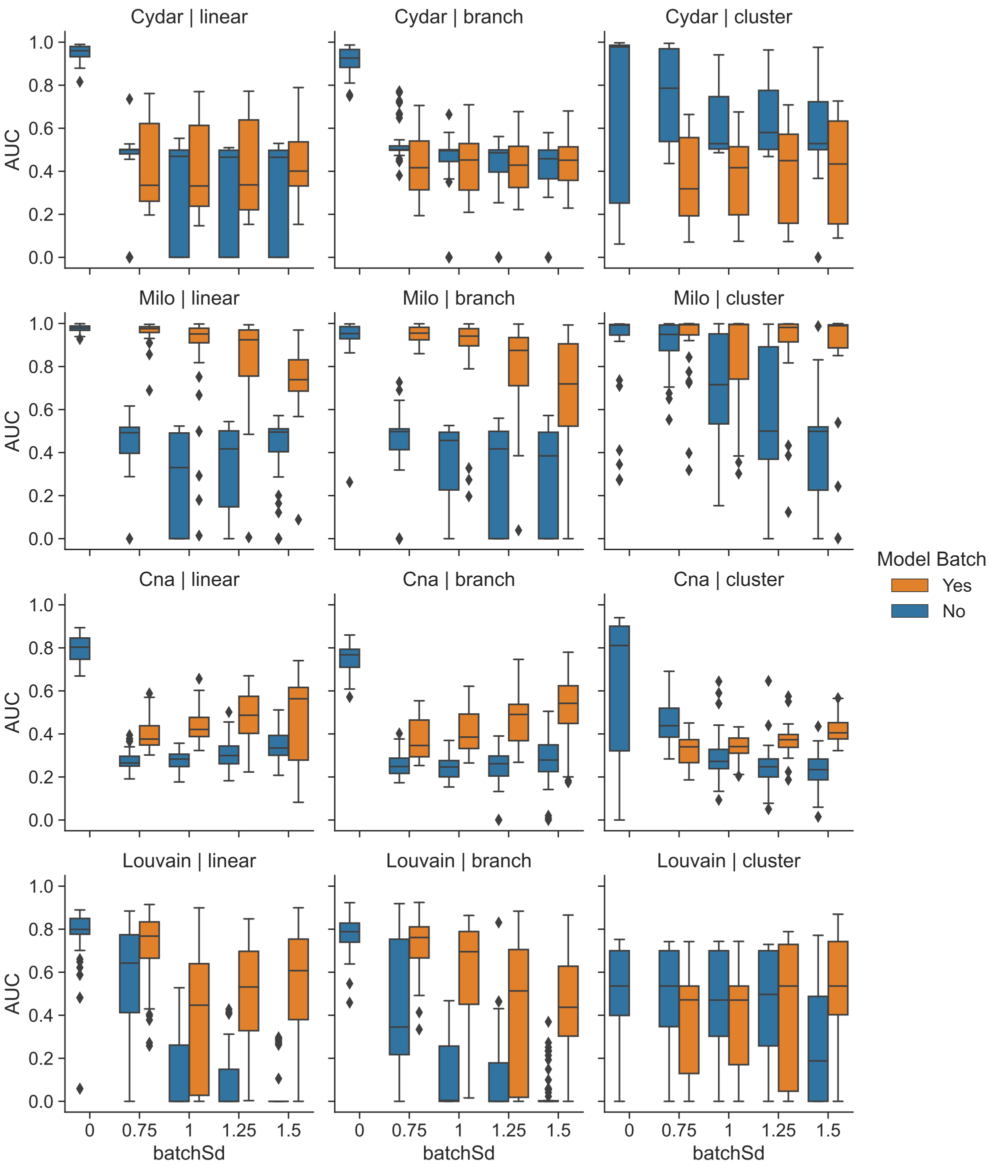


**Figure S3. Performance comparison of DA testing methods by modeling or not modeling batch effects.** Performance of the four DA testing methods (Cydar, Milo, Cna, and Louvain) for DA prediction on the three synthetic datasets (linear (left), branch (middle), and cluster (right)) with batch effects of varying magnitudes (from 0 to 1.5), where the colors (orange and blue) indicate whether batch labels are utilized in the model. The boxplots represent the AUC scores for different target DA cell populations, DA ratios, and random seeds.

#### 2. SUPPLEMENTARY TABLES

**Table S1.** Overview and comparison of DA testing methods.

| Methods | Handle experimental covariates (e.g. B.E.) | Statistical Testing | Approach | DA cells selection | Language |
|---|---|---|---|---|---|
| Cydar [1] | ✔ | ✔ | NB-GLM of HyperSpere | FDR | R |
| Louvain [2] | ✔ | ✔ | NB-GLM of Cluster | FDR | R |
| Milo [3] | ✔ | ✔ | NB-GLM of $k$NN Graph | FDR | R/Python |
| CNA [4] | ✔ | ✔ | SVD of NAM* | FDR | Python |
| DAseq [5] | ✗ | ✔ | Logistic Regression | DA score | R (dep. Python) |
| Meld [6] | ✗ | ✗ | Graph Density Estimation | Likelihood | Python |

*: NAM represents the neighborhood abundance matrix; dep. means depends on.

**Table S2.** Hyperparameters of the six DA testing methods used in the experiments.

| Methods | linear | branch | cluster | COVID-19 PBMC | BCR-XL |
|---|---|---|---|---|---|
| DA-seq [5] | $\mathbf{k} = [30, \ldots, 480]$, where $k_{l+1} - k_l = 50$ . | | | | |
| Meld [6] | $k : 30, \beta : 71$ | $k : 30, \beta : 65$ | $k : 30, \beta : 33$ | $k : 30, \beta : 25$ | $k : 30, \beta : 23$ |
| Milo [3] | $k : 30$ | | | | |
| Cydar [1] | tol : 2.3 | tol : 2.4 | tol : 2.8 | tol : 2.1 | tol : 0.75 |
| Cna [4] | $k : 30$ | | | | |
| Louvain [2] | $k$ : 30, res : 1 | $k$ : 30, res : 1 | $k$ : 30, res : 0.2 | $k$ : 30, res : 0.5 | $k$ : 30, res : 0.6 |

**Table S3.** The corresponding median AUC scores in Figure 3 for the six DA testing methods on the three synthetic datasets (linear, branch, and cluster) with a range of DA ratios (0.75, 0.85, and 0.95).

| | branch | | | cluster | | | linear | | |
|---|---|---|---|---|---|---|---|---|---|
| | 0.75 | 0.85 | 0.95 | 0.75 | 0.85 | 0.95 | 0.75 | 0.85 | 0.95 |
| DA-seq [5] | 0.815 | 0.917 | **0.965** | 0.992 | 0.999 | 0.999 | 0.816 | 0.934 | 0.972 |
| Meld [6] | **0.935** | 0.950 | 0.957 | **1.000** | **0.999** | **0.999** | **0.979** | 0.977 | 0.974 |
| Milo [3] | 0.930 | **0.962** | 0.964 | 0.990 | 0.989 | 0.994 | 0.974 | **0.982** | **0.981** |
| Cydar [1] | 0.919 | 0.928 | 0.933 | 0.975 | 0.981 | 0.978 | 0.941 | 0.965 | 0.962 |
| Cna [4] | 0.702 | 0.780 | 0.807 | 0.774 | 0.890 | 0.857 | 0.727 | 0.807 | 0.849 |
| Louvain [2] | 0.778 | 0.789 | 0.793 | 0.471 | 0.536 | 0.536 | 0.802 | 0.794 | 0.811 |

**Table S4.** The corresponding median AUC scores in Figure 4B for the six DA testing methods on the COVID-19 PBMC and BCR-XL datasets with a range of DA ratios (0.75, 0.85, and 0.95).

| | BCR-XL | | | COVID-19 PBMC | | |
|---|---|---|---|---|---|---|
| | 0.75 | 0.85 | 0.95 | 0.75 | 0.85 | 0.95 |
| DA-seq [5] | 0.773 | 0.896 | 0.956 | 0.802 | 0.882 | 0.935 |
| Meld [6] | **0.978** | **0.976** | **0.972** | **0.954** | **0.957** | **0.962** |
| Milo [3] | 0.902 | 0.940 | 0.959 | 0.838 | 0.891 | 0.924 |
| Cydar [1] | 0.701 | 0.752 | 0.755 | 0.602 | 0.601 | 0.605 |
| Cna [4] | 0.651 | 0.716 | 0.757 | 0.524 | 0.601 | 0.660 |
| Louvain [2] | 0.815 | 0.848 | 0.858 | 0.736 | 0.813 | 0.831 |

**Table S5.** Overview of the hyperparameters in the six DA testing methods.

| Method | Parameter description |
|---|---|
| DA-seq [5] | $\mathbf{k} = [k_1, \ldots, k_l]$: a range of $k$ parameters used to construct $k$-nearest neighbor graphs |
| Meld [6] | $k$: the $k$ parameter to build the $k$-nearest neighbor graph<br>$\beta$: the smoothing parameter applied in graph-based density estimation |
| Milo [3] | $k$: the $k$ parameter to build the $k$-nearest neighbor graph |
| Cydar [1] | tol: the scaling parameter that determines the radius of hyperspheres |
| Cna [4] | $k$: the $k$ parameter to build the $k$-nearest neighbor graph |
| Louvain [2] | $k$: the $k$ parameter to build the $k$-nearest neighbor graph<br>res: the resolution parameter in the Louvain clustering |

**Table S6.** Overall suggestions for the usage of DA testing methods

| Case | Methods | | | | | |
|---|---|---|---|---|---|---|
| | DA-seq[5] | Meld[6] | Milo[3] | Cydar[1] | Cna[4] | Louvain[2] |
| w/o technical noise (e.g. batch effects) | ✔ | ✔ | ✔ | ✔ | ✗ | ✗ |
| w/ technical noise (e.g. batch effects) | ✗ | ✗ | ✔ | ✗ | ✗ | ✗ |
| w/ Type I error control | ✗ | ✗ | ✔ | ✔ | ✔ | ✔ |
| robust to hyperparameter | ✔ | ✔ | ✔ | ✗ | ✗ | ✗ |
| scalable to data size | ✗ | ✗ | ✗ | ✗ | ✔ | ✗ |
